# Staphylococcal and Non-typhoidal *Salmonella* infection statuses in non-human mammals: A potential source of zoonoses in the Greater Accra region of Ghana

**DOI:** 10.1101/2023.10.20.563375

**Authors:** Linda Ama Owusuaa Amoah, Evans Paul Kwame Ameade, Benjamin Yeboah-Ofori, Eric Sampane-Donkor, Langbong Bimi

**Affiliations:** Department of Animal Biology and Conservation Science, University of Ghana, Legon, Accra, Ghana; Department of Pharmacognosy and Herbal Medicine, School of Pharmacy and Pharmaceutical Sciences, University for Development Studies, Tamale, Ghana; Department of Medical Microbiology, Korle Bu, University of Ghana, Legon, Accra, Ghana

**Author notes:** Corresponding author**/** (LAOA). These authors contributed equally to this work (Linda Ama Owusuaa Amoah, Benjamin Yeboah-Ofori, Eric Sampane-Donkor, Langbong Bimi). These authors also contributed equally to this work (Linda Ama Owusuaa Amoah, Evans Paul Kwame Ameade, Benjamin Yeboah-Ofori, Langbong Bimi).

**Keywords:** Zoonoses, Non-typhoidal *Salmonella* infections, Staphylococcal infections, Surveillance, bacterial infections

## Abstract

Background

Bacterial zoonoses are readily transmitted from animals to humans and are thrice more likely to lead to emerging or re-emerging diseases. In Ghana, there is a paucity of animal-related bacterial infection surveillance data, significantly affecting how such diseases are accurately targeted for prevention or control. This study sought to investigate the prevalence of two important bacterial infections in some common animals found in two human-dominated landscapes and ascertain if their prevalence was of imminent public health concern. In most Ghanaian communities, dogs, cats and rodents are non-human mammals that are frequently in contact with humans. As such, they were targeted during this cross-sectional study.

**Methods:** Biological samples collected from animals in households and veterinary institutions were processed using molecular techniques targeting *Staphylococcus* and Non-typhoidal *Salmonella* species. Additionally, medical records were sourced from three (3) major health institutions to determine if cases of bacterial zoonoses were of imminent concern.

**Results:** Overall, the prevalence of staphylococcal and Non-typhoidal *Salmonella* infections were 72.5% and 22.8%, respectively. More animals from the urban areas tested positive for Staphylococcal (χ^2^=5.721; *p*=0.017) and Non-typhoidal *Salmonella* (χ^2^=16.151; *p* < 0.001) infections compared to those from the peri-urban areas. The medical records also revealed that relatively higher cases of staphylococcal infections were reported within three years (2018-2020), although no significant differences were observed between the urban and peri-urban areas.

**Conclusion:** The high prevalence of staphylococcal infections in animals and the high number of hospital cases suggest increased exposure to this bacteria and a higher risk of persons residing in these areas to bacterial zoonoses. Data from the study also suggest that rodents are actively and inactively maintaining the cycle of these two bacterial species and as such, a source of concern. Findings underscore the need for active surveillance of bacterial species with zoonotic potential in non-human mammals regularly found in communities, which is fundamental to developing appropriate disease control strategies.

## Introduction

Nearly all the pandemics that occurred over the last decade originated from animals causing a tremendous impact on the health and economies of nations [1, 2]. Therefore, any pandemic to occur in the future is predicted to have its origin in an animal. Several animal species are reservoirs of zoonoses: they maintain infections in nature and are crucially involved in several zoonotic disease emergence [3,4]. Recently, a more significant part of emerging infectious pathogens has been identified as bacterial [5,6], with zoonotic bacteria thrice more likely to lead to emerging diseases than non-zoonotic bacteria [7].

Compared to animal-related bacterial zoonoses, foodborne bacterial zoonoses attracted much attention in the past [8], similar to the situation in Ghana. However, this trend appears to change as bacterial zoonoses caused by animals are also recognised to impact health systems significantly[9]. For instance, companion animals and rodents transmit important bacteria like *Staphylococcus* and non-typhoidal *Salmonella* species to humans through direct and indirect routes [6,10–11], leading to mild or severe disease outcomes.

Dogs and cats are mostly colonised with *Staphylococcus pseudointermedius,* although many cases have shown that they play vital roles in households and veterinary clinic transmissions of *Staphylococcus aureus* [10, 12–13]. For *Salmonella*, all species are zoonotic except for *Salmonella typhi* and the paratyphoid serotypes [14]. Two key species, *Salmonella enterica* serotype Enteritidis and *Salmonella enterica* serotype Typhimurium, are transferred from animals to humans [15]. Many serotypes have been detected in dogs and cats who may be asymptomatic or symptomatic [16]. Rodents are usually infected with serotypes most common in their environment [14].

The close association of humans with companion animals has resulted in animals like dogs and cats sharing more than 60 parasites [17]. Rodents who live near human settlements offer opportunities for cross-species transmission of pathogens they harbour [18]. Hence, they play significant roles in disease transmission in endemic areas and are a significant risk to sustainable health. In addition, they are a source of food for dogs and cats, a situation that sustains the enzootic transmission cycle of infectious diseases among them [9]. In Ghana, there is a paucity of data on animal-related bacterial infections focusing on mammals which share the same environment with humans. Recently, surveillance has been directed at natural reservoirs of zoonotic infections to enhance disease detection [19], which is fundamental to developing disease control strategies hinged on the one health approach to avert any potential epidemic or pandemic.

## Materials and methods

### Study design

This study was cross-sectional. Primary and secondary data were collected from two human-dominated landscapes (landscapes heavily dominated by people) in the Greater Accra Region. The urban and peri-urban areas were the two human-dominated landscapes selected for this study, with data collection spanning from October 2019 to October 2020.

Two non-probability sampling approaches, purposive and convenient sampling techniques, were used in the selection of the study populations. This was because there existed no reliable data on the population of targeted animals in the Greater Accra Region to serve as a sampling frame at the time of this study. Both techniques were used to obtain an eligible and representative sample size of animal subjects for this research.

### Study setting

There is no universally accepted definition of ‘urban’ as most definitions are based on national definitions or characterizations [20] In Ghana, communities with about five thousand (5,000) persons or more are categorised as urban [21]. However, for this research, the use of the minimum population size alone seemed simplistic as over 87% of the population of the Greater Accra region, per this definition is in urban areas [21]. Therefore, the minimum population size and the density of socio-economic activities in such areas were considered in selecting the urban and peri-urban areas from the 29 Metropolitan, Municipal and District assemblies (MMDAs) in the Greater Accra region [22]. Legon and Madina were purposively selected from the Accra Metropolitan Assembly and La-Nkwatanang Madina Municipality, respectively, under the urban category as both study sites are completely urbanised. For this study, Legon comprised the University of Ghana campus (5.6506°N, 0.187°W) and East Legon (5.6358°N, 0.1614°W).

Legon shares close boundaries with Madina (5.6731°N, 0.0980°W), the administrative capital of the La Nkwantanang-Madina Municipal [21]. Both communities are characterised by higher educational institutions and good health facilities. Economically, these two communities are very vibrant, and most of the populations are engaged in commerce.

Dodowa (5.8829° N, 0.00980W) in the Shai-Osudoku District was selected for the peri-urban category because this town was formed due to urban sprawl. Additionally, it exhibits both urban and rural characteristics, as peri-urbanisation connotes.

### Study population and sample size determination

Rodents and companion animals (dogs and cats) were targeted for this study because of the increased interaction or contact of these animals with humans in most Ghanaian communities. Since there is no published data on the prevalence of these two bacterial infections among the targeted animal species, a minimum sample size of 384 was obtained at a 95% confidence interval, assuming a minimum prevalence of 50% using the formula below. This formula, according to Naing et al. [23], is used in calculating sample size which seeks to estimate population prevalence with good precision.

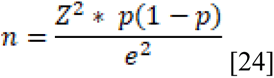

Where n is the sample size, Z represents the Z score at a 95% confidence interval of 1.96; *p* is the minimum estimated prevalence (50%); e=margin of error/absolute error (5%) at a 95% confidence interval.

In all, a minimum of 384 animals (dogs, cats and rodents) were to be sampled using a purposive sampling technique due to the absence of a sampling frame as previously explained.

### Sources of data

Biological samples were collected from targeted animals from households, veterinary institutions, and around human settlements in selected study settings after ethical clearance and informed consent were given. In the urban areas, two veterinary institutions were selected: the University of Ghana Small Animal Teaching Hospital (UG-SATH) and the East Legon Veterinary Centre (ELVC). Four (4) criteria were used for the selection of these veterinary institutions: location, accessibility, range of services provided and patronage. Unfortunately, there was no functional veterinary clinic in Dodowa at the time of data collection.

Medical records from 2018-2020 on *Staphylococcus* and Non-typhoidal *Salmonella* infections were sourced from three (3) major health institutions: the University of Ghana and Pentecost Hospitals (urban category) and the Shai-Osudoku District Hospital (peri-urban category). The purpose of these secondary data was to corroborate or contradict the possibility of bacterial zoonotic disease transmission in the study areas.

These three hospitals were also selected using four (4) criteria: accessibility, range of services provided, availability of requisite facilities and patronage. Even though there are a few other health institutions in the study areas, most could not meet these criteria.

The University of Ghana Hospital, referred to as the Legon Hospital is a quasi-government hospital that offers both general services and specialist clinics. It has a 130-bed capacity which consists of a paediatric unit, maternity, casualty and emergency ward. The Pentecost Hospital, situated in Madina is a mission hospital. It is the municipal hospital for the La-Nkwantanang Madina municipality and provides services in obstetrics and Gynaecology, child health, accident and emergency services as well as other relevant services [25]. Comparatively, this hospital has relatively higher patronage. The third hospital, the Shai-Osudoku District Hospital located in Dodowa, is the only major hospital in the Shai-Osudoku district with advanced medical facilities. The hospital offers a variety of clinical services to people from all spheres of life in the district and beyond.

### Sample collection

#### Rodent sampling

Four sampling sites in each human-dominated landscape were used for trapping rodents. The sampling areas had an average of 50 trap lines at least 10 metres apart. A small amount of bait (a mixture of peanut butter, palm nut and maise flour) was placed in each trap which was then labelled and placed under logs near burrows, homes, farmlands, and dumpsites. The traps were left overnight and inspected daily for three consecutive days. After each daily inspection, the traps were re-baited when necessary. Active traps (traps containing rodents) were recorded and carefully handled according to standard protocols [26]. The rodents were described using morphometric keys. Any trap found missing, sprung, or not triggered was regarded as a misfire (capture of non-target species) and was also recorded.

#### Companion animals sampling

Dogs and cats were sampled from households and veterinary institutions at Legon-Madina and Dodowa after their owners consented to participate in the study.

#### Collection of blood samples from animals

Less than 1 ml of venous blood was collected from the animals into heparinised tubes adhering to standard protocol and procedures for blood collection and transportation. Blood was collected from the tail region of rodents and the cephalic veins of dogs and cats by a veterinary officer adhering to sterile safety measures.

On some occasions, swabbing the cloaca with sterile cotton was employed when animals exhibited aggression during blood collection. The swabs were placed into appropriately labelled tubes and transported to the laboratory under cold chain conditions.

### Processing of biological samples

Deoxyribonucleic acid (DNA) of pathogens was extracted from blood and swabs using the protocol described by Soumet et al. [27] with modifications. A total of 100 µL blood was inoculated in 900 μL Luria Bertani broth (Invitrogen 12780-052), and the culture was incubated at 37℃ for 48 h in a shaking incubator (9.44 x 10^-2^ xg). Afterwards, 1 ml of individual culture was picked and centrifuged for three (3) min at 13000 xg (4℃), after which the resultant pellets were washed twice in 500 μL 1X TAE buffer (Tris, Acetic Acid, and EDTA; pH 8.0), and centrifuged at 13000 xg for 3 min (4℃). The pellets were re-suspended in 200 μL sterile water, boiled for 8 min, and then stored at -20℃.

The swabs were eluted in 1000 μL of Phosphate Buffered Saline (PBS) after vortexing for 2 min before extracting the bacterial DNA, as explained earlier.

### DNA Amplification by Polymerase Chain Reaction

Direct Polymerase Chain Reaction (PCR) was used to amplify numerous copies of diminutive DNA fragments using genus-specific primers. Further details of the primer set used for the PCR are listed in Table 1.

**Table 1.**
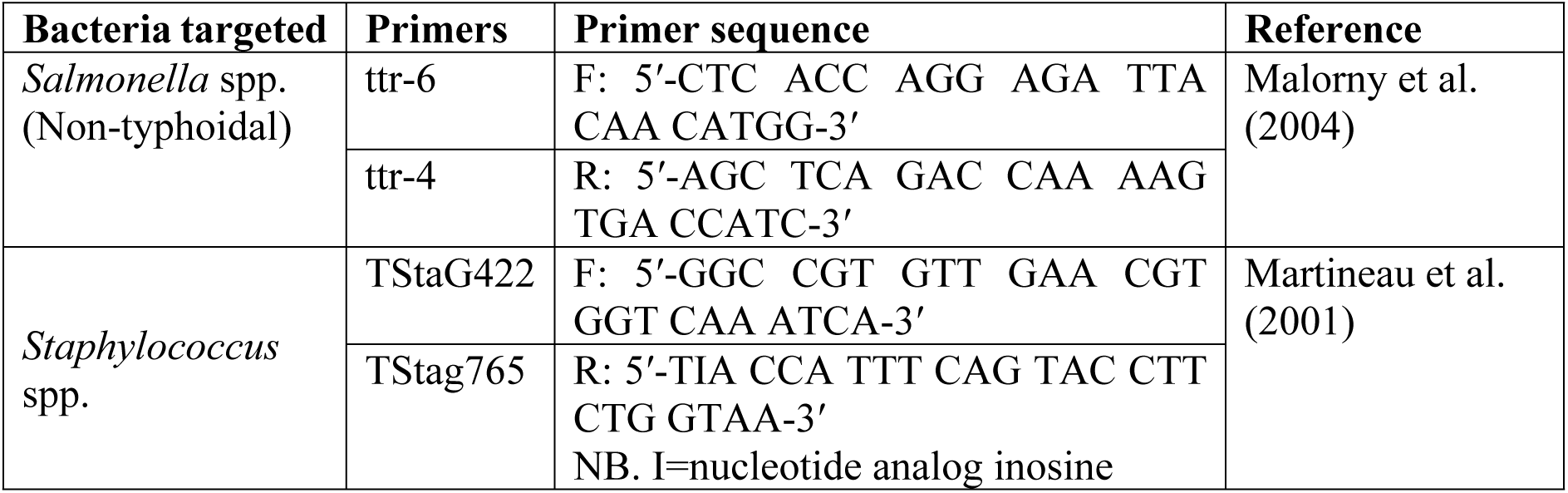
Primer sets for the polymerase chain reaction (PCR)

All PCR reactions were performed in final volumes of 25.0µL containing 7.50µL sterile nuclease-free PCR water, 12.50µL OneTaq^®^ Quick-Load^®^ 2X Master Mix with Standard Buffer (New England Biolabs Inc.), 0.5µL each of forward and reverse primers and 4.0 µL DNA template. This was done at initial and final denaturation of 95℃ for 2min and 94℃ for 40s, respectively. The annealing temperature was set at 53℃ for 1 min, followed by an initial extension at 72℃ for 3 min. The final extension was at 72℃ for 7 min with a holding temperature of 4℃. The PCR products were examined by electrophoresis on 1.2% agarose gels stained with 4.0 µL Ethidium bromide. The wells were each loaded with 6.0 µL PCR product and run at 80V for 90 min. Fluorescence from the DNA-bound Ethidium bromide was then visualised under Ultra Violet (UV) Trans-illuminator (SYNGENE Inc.). The amplified products were measured using a 100 base pair (bp) DNA ladder (New England Biolabs Inc.) at specific bands corresponding to their base pairs for the two bacterial species.

### Analysis of data

Data were entered into Microsoft Excel (2006) and subsequently analysed with IBM Statistical Package for Social Sciences (SPSS version 20). Comparisons were made using the Mann-Whitney U test, Student’s t-test, Pearson Chi-square or Analysis of variance (ANOVA) depending on the nature of the distribution of the data (normal or non-normal distribution). The prevalence of infection was also calculated using the formula below

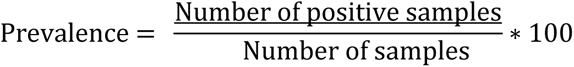

At a 95% confidence level, *p*<0.05 was considered statistically significant.

### Ethical approval and informed consent

This study (UG-IACUC 009/18-19) was approved by the University of Ghana Institutional Animal Care and Use Committee (UG-IACUC). Ethical clearance or approval was also obtained from selected veterinary and health institutions as well as the Ghana Health Service. Informed consent was also sought from owners of companion animals before their animals were sampled.

### Results

### Distribution of animal species

A total of 404 biological samples were collected from dogs (n=185, 45.8%), cats (n=15,3.7%) and rodents (n=204, 50.5%) (Table 2). In all, fewer cats were sampled than dogs from households and veterinary institutions as shown in Table 2.

**Table 2.**
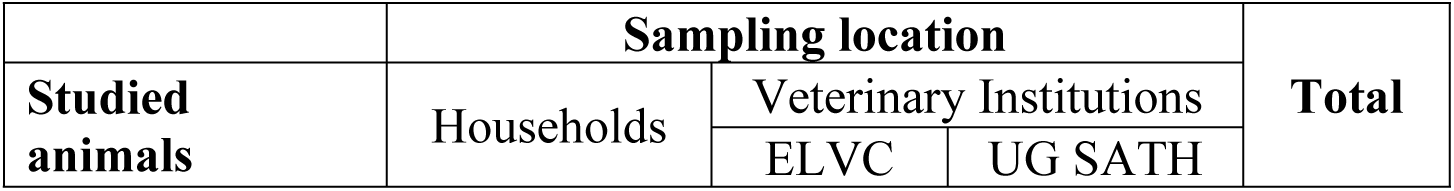
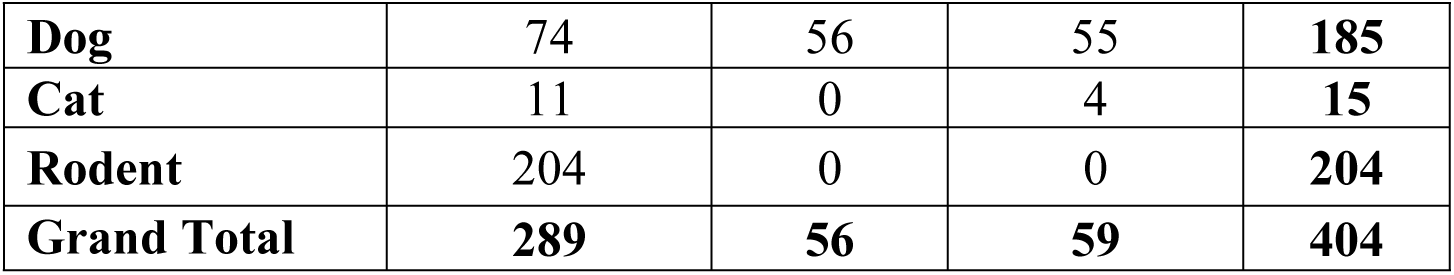
Distribution of sampled animals in the two sampling locations.

A total of one-hundred and fifteen (115) companion animals consisting of 111 (96.52%) dogs and 4(3.48%) cats were sampled from the two veterinary institutions in the urban areas.

For the rodents, 204 individuals belonging to seven (7) species were trapped. There were variations in the distribution of trapped rodents in the two human-dominated landscapes (Table 3). Statistically, significant differences were found between their distributions in the urban and peri-urban areas using the Mann-Whitney U test (U=2770.00, Z=-4.349, *p*<0.001) (S1 Table). *Arvicanthis niloticus* was the dominant species trapped in the urban communities in contrast to *Praomys tullbergi,* the dominant species from the peri-urban areas. However, overall, *A. niloticus* was the dominant rodent, and the least was *Crocidura olivieri* (Table 3).

**Table 3.** Distribution of rodents in the human-dominated landscape.

| **Rodent** | **Human-Dominated Landscape** |  | **Total** |
| --- | --- | --- | --- |
|  | Urban | Peri-Urban |  |
| *Praomys tullbergi* | 0<br>(0.0%) | 38<br>(100.0%) | 38<br>(100.0%) |
| *Arvicanthis niloticus* | 113<br>(100.0%) | 0<br>(0.0%) | 113<br>(100.0%) |
| *Rattus rattus* | 9<br>(47.4%) | 10<br>(52.6%) | 19<br>(100.0%) |
| *Cricetomys gambianus* | 5<br>(100.0%) | 0<br>(0.0%) | 5<br>(100.0%) |
| *Rattus norvegicus* | 18<br>(100.0%) | 0<br>(0.0%) | 18<br>(100.0%) |
| *Mus musculus* | 0<br>(0.0%) | 10<br>(100.0%) | 10<br>(100.0%) |
| *Crocidura olivieri* | 0<br>(0.0%) | 1<br>(100.0%) | 1<br>(100.0%) |
| **$U=2770.00$, \* $p<0.001$** |  |  |  |

The asterisk (\*) represents a statistically significant result.

### Distribution of animals in the two human-dominated landscapes

Out of the total number sampled, 81.08% of the dogs, 86.67% of cats and 71.08% of rodents were from urban areas (S2 Table). More animals were sampled from the urban areas compared to the peri-urban areas. This was confirmed using the Mann-Whitney *U* Test, which was statistically significant (*U*=12619.500, *p*=0.019)(S3 Table).

The distribution of the sampled animals from the two human-dominated landscapes is illustrated in Fig 1.

**Fig 1.**
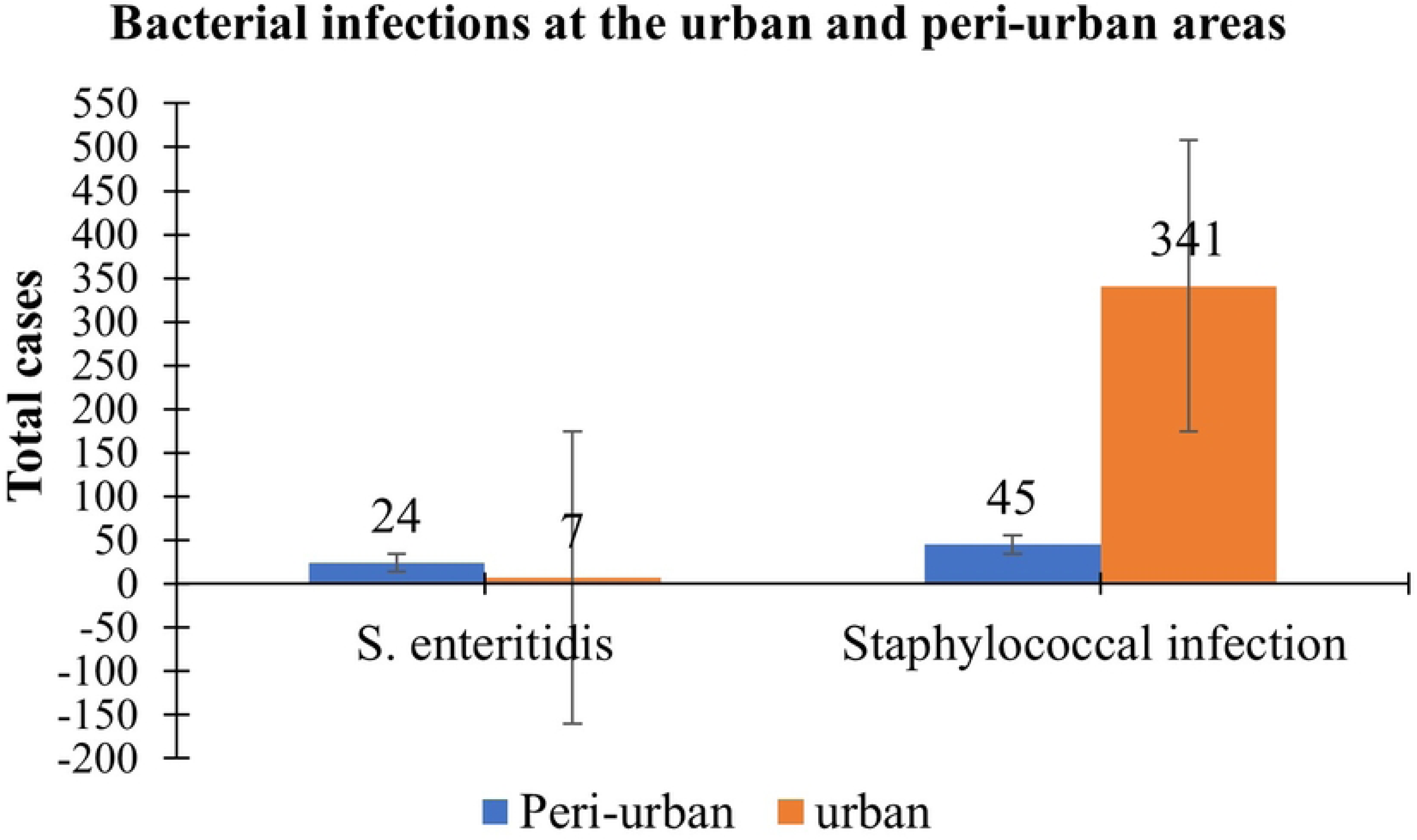
Distribution of sampled animals in the two human-dominated landscapes.

### Prevalence of bacterial infections

Results from the PCR showed more animals, 72.5% (95% CI= 67.8%-76.8%) were positive for *Staphylococcus* spp. compared to Non-typhoidal *Salmonella* spp., 22.8% (95% CI=18.8%-27.2%). Fig 2 presents the prevalence of Staphylococcal and Non-typhoidal *Salmonella* infections in the targeted animals.

**Fig 2.**
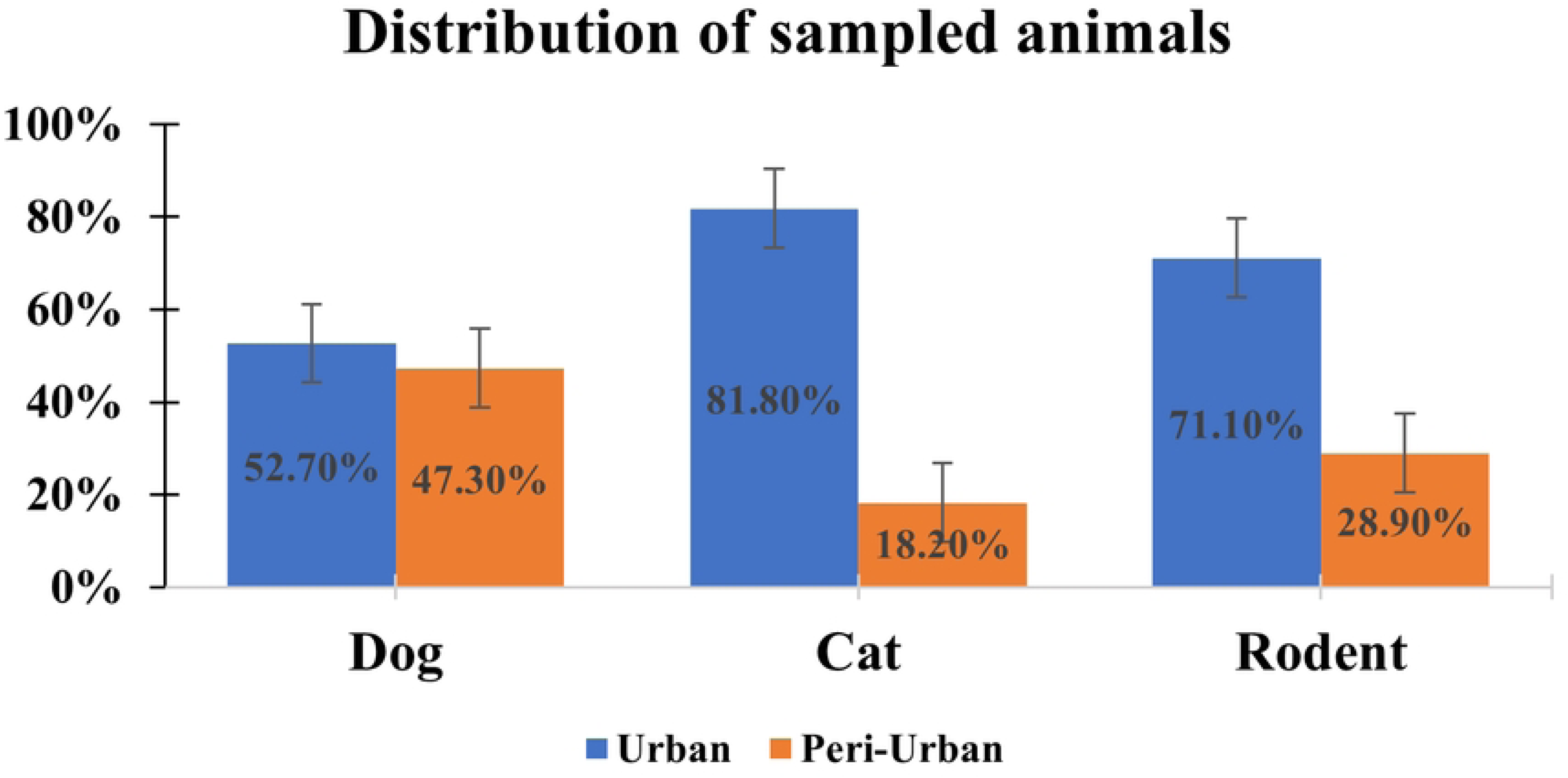
Prevalence of bacterial infections among sampled animal species.

### Bacterial infections in the two human-dominated landscapes

Human-dominated landscapes are critical loci for disease transmission, and therefore prevalence was analysed in relation to the urban and peri-urban areas. A Pearson Chi-square analysis showed a high number of the sampled animals tested positive for both bacteria species in the urban areas in contrast to the peri-urban areas (Table 4).

**Table 4.** Prevalence of bacterial infections in the two human-dominated landscapes.

| Bacteria |  | Human-dominated landscape |  | Total | Pearson Chi-square |
| --- | --- | --- | --- | --- | --- |
|  |  | Urban | Peri-urban |  |  |
| *Salmonella* spp. (Non-typhoidal) | Negative | 253<br>(81.1%) | 59<br>(18.9%) | 312<br>(100.0%) | $\chi^2=16.151$<br>$*p < 0.001$ |
|  | Positive | 56<br>(60.9%) | 36<br>(39.1%) | 92<br>(100.0%) |  |
| *Staphylococcus* spp. | Negative | 94<br>(84.7%) | 17<br>(15.3%) | 111<br>(100.0%) | $\chi^2=5.721$<br>*$p=0.017$ |
|  | Positive | 215<br>(73.4%) | 78<br>(26.6%) | 293<br>(100.0%) |  |

The asterisk (\*) represents a statistically significant result.

### Bacterial infections among the sampled animals

In comparing infection prevalence among the three sampled animals, the highest prevalence of Staphylococcal and Non-typhoidal *Salmonella* infections were found in rodents. However, in dogs, the prevalence was also high. A Pearson Chi-square analysis revealed that these differences were not statistically significant (Table 5).

**Table 5.** Prevalence of bacterial infections among the sampled animal species.

| Bacteria | Infection status | Sampled animals |  |  | Total | Pearson Chi-square |
| --- | --- | --- | --- | --- | --- | --- |
|  |  | Dog | Cat | Rodent |  |  |
| *Staphylococcus* spp. | Negative | 48<br>43.2% | 5<br>4.5% | 58<br>52.3% | 111<br>100.0% | $\chi^2=0.569$<br>$p=0.752$ |
|  | Positive | 137<br>46.8% | 10<br>3.4% | 146<br>49.8% | 293<br>100.0% |  |
| Total |  | 185<br>45.8% | 15<br>3.7% | 204<br>50.5% | 404<br>100.0% |  |
| *Salmonella* spp. (Non-typhoidal) | Negative | 144<br>46.2% | 9<br>2.9% | 159<br>51.0% | 312<br>100.0% | $\chi^2=2.630$<br>$p=0.269$ |
|  | Positive | 41<br>44.6% | 6<br>6.5% | 45<br>48.9% | 92<br>100.0% |  |
| Total |  | 185<br>45.8% | 15<br>3.7% | 204<br>50.5% | 404<br>100.0% |  |

Although higher infection prevalence was recorded from animals sampled from the households compared to the veterinary institutions, no statistically significant differences were observed (Table 6).

**Table 6.** Prevalence of bacterial infections in animals from households and veterinary institutions.

| Bacteria |  | Sampling location |  | Total | Pearson Chi-square |
| --- | --- | --- | --- | --- | --- |
|  |  | Household | Veterinary Institutions |  |  |
| *Salmonella* spp. (Non-typhoidal) | Negative | 225<br>72.1% | 87<br>27.9% | 312<br>100.0% | $\chi^2=0.227$<br>$p=0.634$ |
|  | Positive | 64<br>69.6% | 28<br>30.4% | 92<br>100.0% |  |
| **Total** |  | 289<br>71.5% | 115<br>28.5% | 404<br>100.0% |  |
| *Staphylococcus*<br>spp. | Negative | 83<br>74.8% | 28<br>25.2% | 111<br>100.0% | $\chi^2=0.789$<br>$p=0.374$ |
|  | Positive | 206<br>70.3% | 87<br>29.7% | 293<br>100.0% |  |
| **Total** |  | 289<br>71.5% | 115<br>28.5% | 404<br>100.0% |  |

### Images of gel electrophoresis

PCR products were analysed using agarose gel electrophoreses, and the images for Non-typhoidal *Salmonella* and *Staphylococcus* spp. are shown in Figs 3 and 4. The molecular weight marker, or the DNA ladder denoted by M (Lane M), was used to detect the approximate sizes of the amplicons.

**Fig 3.**
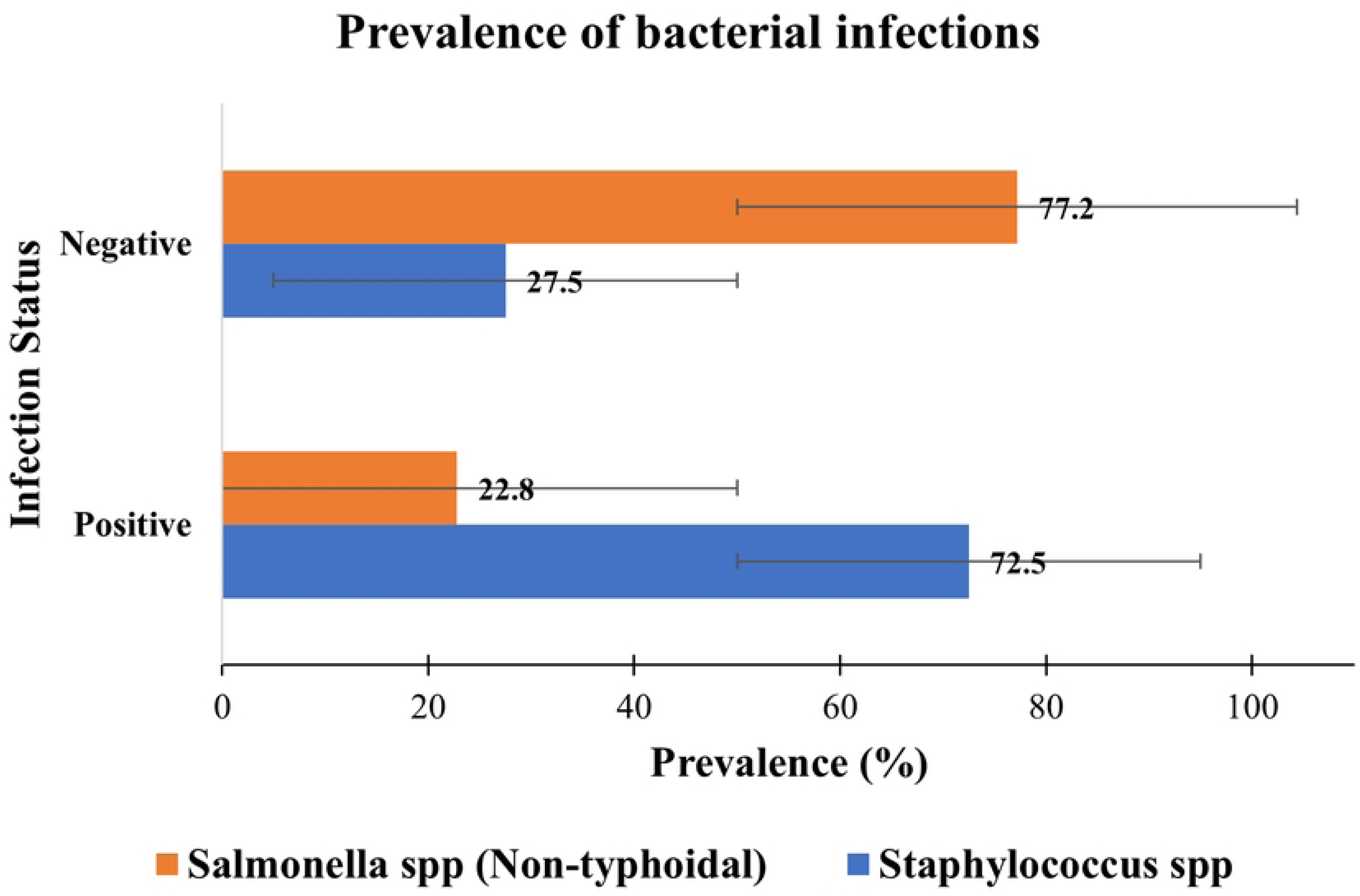
Amplified PCR product of *Salmonella* spp. (Non-typhoidal) (Lane M=100 bp, Lane 1-15, 17-26=positive, Lane 16=negative) Lanes 1-26 contain DNA from the blood of sampled animals after preparation with DNAzol and PCR. Gel depicting 350 bp PCR product.

**Fig 4.**
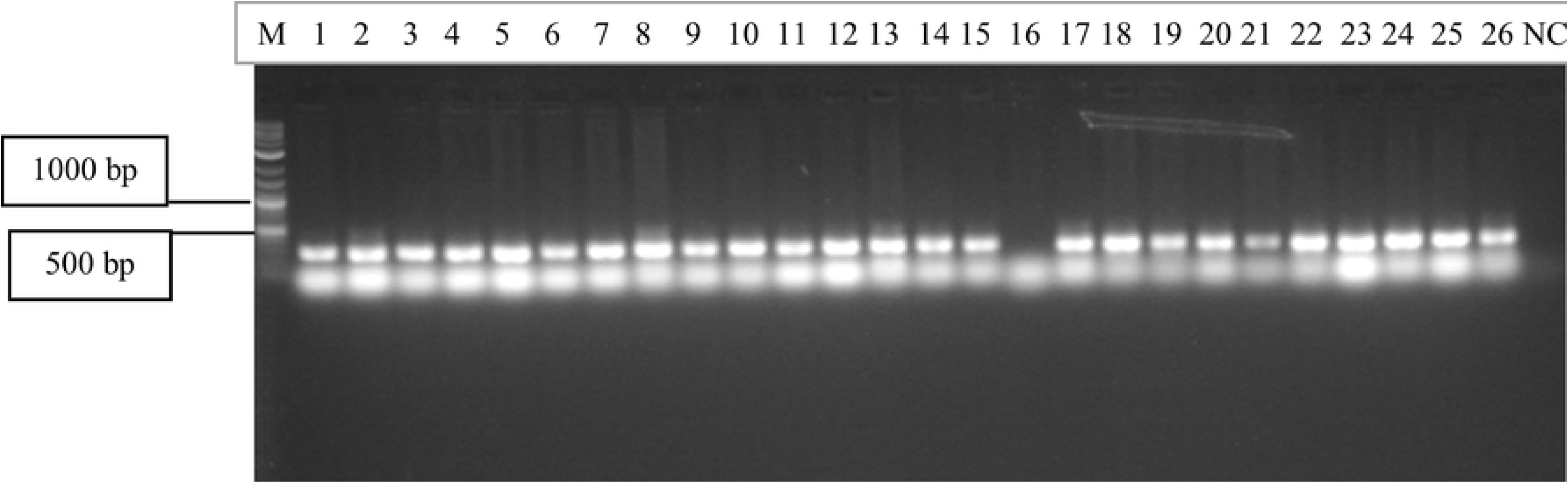
Amplified PCR product of *Staphylococcu*s spp. Lane M=100 bp, Lane NC= negative control, Lanes 10, 11, 13, 16 and 18= positive, Lane 1-9, 12,14-15, 17, 19=negative) Lanes 1-19 contain DNA from the blood of sampled animals after preparation with DNAzol and PCR. Gel depicting a 550 bp product.

### Medical records from hospitals in the two human-dominated landscapes

Secondary data from two out of the three hospitals revealed cases of bacterial zoonotic infections. In scrutinising bacterial zoonoses presented at the hospitals, bacterial infections that were non-zoonotic (S4 Table) were excluded. Therefore, cases of zoonotic bacterial infections and bacterial infections with zoonotic potential presented by clients at the two hospitals are shown in Table 7.

**Table 7.**
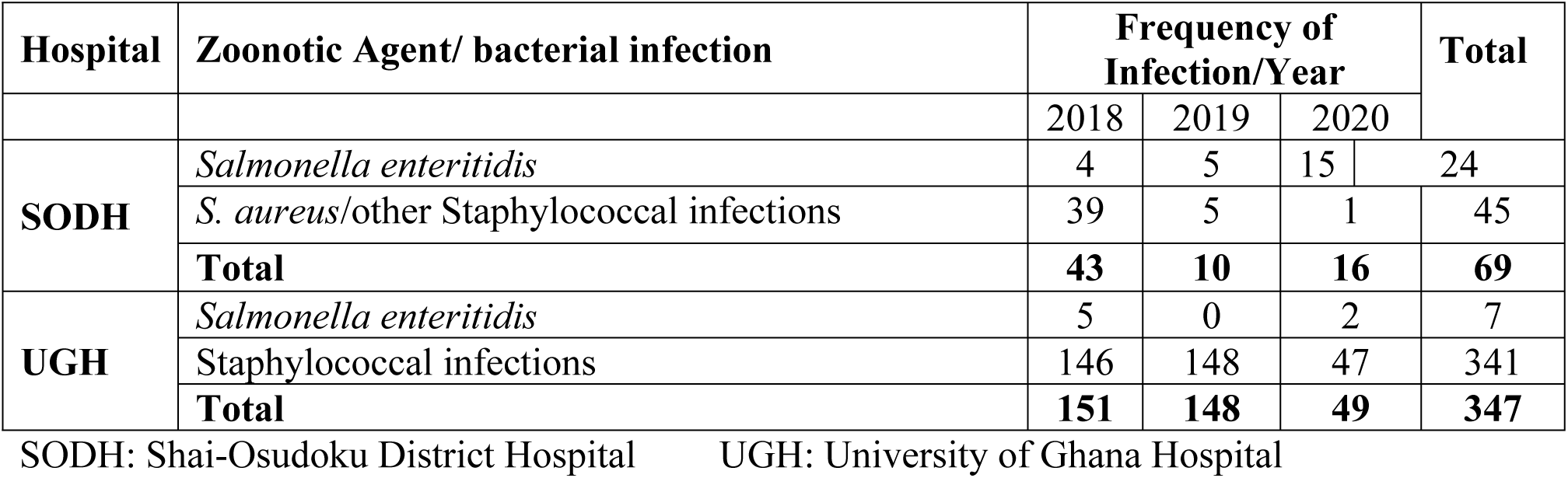
Bacterial infections presented by clients at the hospitals.

In comparing cases of Non-typhoidal *Salmonella* to staphylococcal infections, both hospitals reported relatively higher cases of staphylococcal infections. Nevertheless, there were no statistically significant differences (*p*>0.05) in these two bacterial infections between the urban and peri-urban areas(S7 Table).

In addition, records of staphylococcal infections from SODH showed a decline from 2018 to 2020, whereas that of *Salmonella enteritidis* increased. On the other hand, Staphylococcal infections were very high in UGH until the number of cases decreased considerably in 2020. On the whole, cases of bacterial zoonoses recorded in UGH were the lowest in 2020 compared to SODH. However, there were no significant differences between the two bacterial infections recorded over the three years (*p*>0.05)(S8 Table).

Gender-based records of the two bacterial infections presented at the hospitals were also examined and the results shown in Table 7.

**Table 7.** Gender differences in bacterial infections presented at the hospitals.

| Hospital | Zoonotic agent/ bacterial infection | 2018 |  | 2019 |  | 2020 |  |
| --- | --- | --- | --- | --- | --- | --- | --- |
|  |  | Male | Female | Male | Female | Male | Female |
| SODH | *S. enteritidis* | 3 | 1 | 4 | 1 | 6 | 9 |
|  | *S. aureus*, other staphylococcal infection | 21 | 18 | 2 | 3 | 1 | 0 |
| U. G | *S. enteritidis* | 3 | 2 | 0 | 0 | 2 | 0 |
|  | Staphylococcal | 72 | 74 | 63 | 85 | 20 | 27 |
|  | infection |  |  |  |  |  |  |
| **Total** |  | 99 | 95 | 69 | 89 | 29 | 36 |

SODH: Shai-Osudoku District Hospital UGH: University of Ghana Hospital

Records of these infections between males and females were also compared and the results showed that on average, more females (M=17.416, SD=27.73) than males (M=16.42, SD=24.93) were diagnosed with these bacterial infections (S9 Table). Even so, an independent t-test showed that the difference was not statistically significant, t (22) =-0.093, *p*=0.927 (S9 Table).

Again, a cursory look at the total cases of these two bacterial infections reported at the hospitals appeared to differ between the urban and peri-urban areas as illustrated in Fig 5.

**Fig 5.**
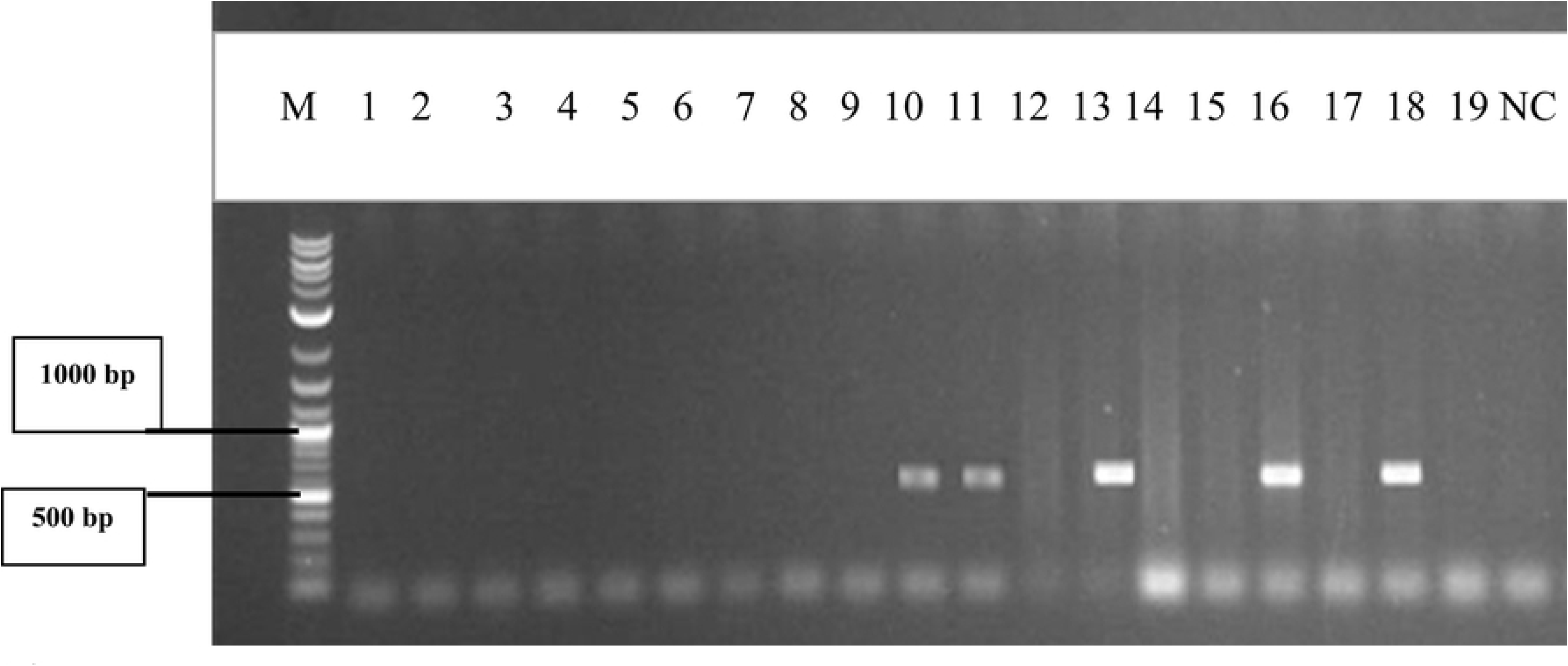
Bacterial infections in the two human-dominated landscapes.

An independent t-test confirmed there existed a significant difference in the total cases of *S. enteritidis* and Staphylococcal infections reported in the urban (M=28.08, SD=32.73) and peri-urban areas (M=5.75, SD=6.92).

Further comparison was also done between the type of bacterial infection and the total cases recorded. The medical records showed that more staphylococcal infections (M=31.25, SD=30.59) were reported than *S. enteritidis* (M=2.583, SD=2.712) (S11 Table). Using the independent t-test, the difference was found to be statistically significant, t (11.173) =-3.233, *p*=0.008 (S11 Table).

## Discussion

Bacterial zoonoses are readily transmitted from animals to humans, and reports suggest that a higher number of zoonoses are caused by bacterial agents [6]. Therefore, this study aimed to ascertain if the prevalence of these two important bacterial species was of imminent concern in the Greater Accra region. As dogs, cats and rodents are non-human mammals frequently found in most communities in Ghana [28] they were targeted during this study. The prevalence of *Staphylococcu*s and Non-typhoidal *Salmonella* infections were 72.5% and 22.8 %, respectively.

Of the three animals examined, rodents recorded the highest infection prevalence. Rodents are known natural hosts of *Staphylococcus* spp. [29]. Thus, many rodents that tested positive for *Staphylococcus* spp. were not out of place. However, this data indicates that rodents around human settlements significantly increase the risk of zoonotic bacterial infections due to high exposure. Besides, transmission may be indirect when companion animals (dogs and cats) feed on these rodents, get infected and consequently transmit these infections to their owners [30]. This high prevalence of *Staphylococcus* spp. in rodents corroborates that of Ribas et al. [12] in Thailand. In comparison to other works, the prevalence of *Staphylococcus* spp. among dogs was higher (46.8%) than what was reported by Han et al. [31] in South Korea and Qekwana et al. [32] in South Africa, where a prevalence of 37% and 27% respectively, were recorded.

Additionally, the most dominant rodent trapped, *Arvicanthis niloticus* is identified as an important agricultural pest with extensive distributions in Ghana and Africa. This implies that high populations of these rodents could cause significant pre- and post-harvest losses of agricultural products and increase the probability of human exposure to the diseases they transmit [28].

Research has shown that *Salmonellae* are commonly distributed by domestic and wild animals that maintain the animal-to-animal cycle through the usual faecal-oral route [15]. The high prevalence of dogs and rodents being positive suggests the environment may be contaminated with this bacterium because these animals get infected by ingesting infected faeces from the environment. Subsequently, the shedding of the parasite by dogs, cats and rodents may lead to the potential transmission of zoonotic *Salmonella* infections to humans in the same household. Though dogs and cats are usually subclinical, Marks et al. [33] believed that infections could progress from mild to fatal conditions of gastroenteritis and septicaemia. Therefore, the importance of this relatively high infection prevalence, particularly in dogs, cannot be overemphasised. In this study, the prevalence of *Salmonella* spp. in dogs was relatively higher than what was reported by Núñez Castro et al. [34] (6.27%) and Bataller et al. [35] (1.85%). The differences could be because both studies sampled healthy dogs, whereas, in this study, both healthy and unhealthy dogs were sampled.

Even though positive cases of Staphylococcal and *Salmonella* infections from hospitals within the study areas were low, there is an indication of possible zoonotic bacterial transmissions in urban and peri-urban communities. Data from the urban and peri-urban categories revealed a high record of staphylococcal infections, consistent with prevalence from the sampled animals. The high cases of staphylococcal infections from rodents and dogs suggest that persons living in such areas are at a higher risk of zoonotic staphylococcal diseases due to the high exposure.

Findings from the study demonstrated that the distribution of the various rodent genera between the two human-dominated landscapes was statistically significant, indicating that some rodents thrived better in one area than the other. This data corroborates the study by Assefa & Chelmala [36], where the distribution and diversity of rodents were found to differ across habitats. The two dominant rodents, *Arvicanthis niloticus* and *Praomys tulbergi,* found in the urban and peri-urban areas, respectively, suggest that they might be the most typical species found in these areas which merit further surveillance in terms of zoonotic disease transmission.

Furthermore, the high abundance of rodents recorded from the urban areas in contrast to the peri-urban areas which have a combination of rural and urban characteristics and good vegetation is worth investigating. This could probably be due to poor sanitary conditions such as the irregular collection of garbage and open sewers that often characterise urban spaces in the Greater Accra Region. In the view of Panti-May et al. [37], such conditions create favourable conditions for rodents to thrive. On the other hand, one reason for the low numbers of rodents in the peri-urban areas may be the continuous rodent-hunting activities in these areas, as observed during the data collection.

## Conclusion

The high prevalence of staphylococcal infections in animals and the high number of hospital cases suggest increased exposure to this bacteria and a higher risk of persons residing in these areas, especially urban communities, to bacterial zoonoses. Results from this study indicate that rodents are actively and inactively maintaining the cycle of these two bacterial species which could threaten the sustainable health of persons found in those areas. Furthermore, the data shows that these two bacteria merit continued investigations in our urban and peri-urban areas. Lastly, the findings underscore the need for active surveillance of zoonotic and possible zoonotic bacterial diseases in non-human mammals regularly found in our communities, which is fundamental to developing zoonotic disease control programmes using the one health approach.

## Acknowledgements

We wish to thank all the veterinary and health institutions that participated in this study. Special thanks to the University of Ghana-BANGA-Africa Project for their immense support.

## Supporting information

S1 Table. Comparison of the distribution of rodents between the two human-dominated landscapes

S2 Table. Distribution of animals in the two human-dominated landscapes

S3 Table. Comparison of the distribution of animals in the two human-dominated landscapes

S4 Table. Summary of the two bacterial infections documented by the hospitals from 2018 to 2020

S5 Table. Comparison of staphylococcal infections between the two hospitals

S6 Table. Comparison of *Salmonella* (non-typhoidal) infections between the two hospitals

S7 Table. Comparison of bacterial infections recorded by the hospitals from 2018-2020

S8 Table. Comparison of bacterial infections recorded by the hospitals in the human-dominated landscapes.

S9 Table. Comparison of total cases of bacterial infections between males and females

S10 Table. Comparing total cases between the two human-dominated landscapes

S11 Table. Differences in the total number of cases reported for the two bacterial infections

## Notes

### Competing Interest Statement

The authors have declared no competing interest.

